# Neural and computational evidence for a predictive learning account of the testing effect

**DOI:** 10.1101/2025.03.17.643739

**Authors:** Haopeng Chen, Pieter Verbeke, Stefania Mattioni, Cristian Buc Calderon, Tom Verguts

**Affiliations:** Department of Experimental Psychology, Faculty of Psychology and Educational Sciences, Ghent University, Ghent, 9000, Belgium; AI Lab, Howest University of Applied Sciences, Kortrijk, 8500, Belgium; Centro Nacional de Inteligencia Artificial (CENIA), Santiago, 8320000, Chile

**Keywords:** Testing effect, Predictive learning, fMRI, Ventral Striatum, Insula

## Abstract

Testing enhances memory more than studying. Although numerous studies have demonstrated the robustness of this classic effect, its neural and computational origin remains debated. Predictive learning is a potential mechanism behind this phenomenon: Because predictions and prediction errors (mismatch between predictions and feedback) are more likely to be generated in testing (relative to in studying), testing can benefit more from predictive learning. We shed light on the testing effect from a multi-level analysis perspective via a combination of cognitive neuroscience experiments (fMRI) and computational modeling. Behaviorally and computationally, only a model incorporating predictive learning can account for the full breadth of behavioral patterns and the robust testing effect. At the neural level, testing and prediction error both activate the canonical reward-related brain areas in the ventral striatum, insula, and midbrain. Crucially, back sorting analysis revealed that activation in the ventral striatum, insula, and midbrain can enhance declarative memory. These results provide strong and converging evidence for a predictive learning account of the testing effect.

**Significance Statement:** An exam is not a neutral measurement of memory. The *testing effect* entails that a test (e.g., an exam), is more effective than study for learning and memory. The same effect can be harnessed also before events of significance take place, rendering it an important aspect of an active learning strategy. Nevertheless, its origin remains unknown. We propose a novel predictive learning account, which posits that testing (vs studying) facilitates predictions about study material and promotes learning from prediction errors. Computationally, the testing effect was explained through a predictive-learning-based neural network. Neurally, testing and prediction error activate common neural areas, which in turn enhance declarative memory. This account may extend beyond testing to support active learning.

## Introduction

A remarkable finding is the testing effect, the robust phenomenon that testing enhances declarative memory retention more effectively than studying (1, 2). For instance, after an initial study session, taking tests instead of engaging in repeated study helps one to retain knowledge (3–5). Accordingly, learning apps such as Duolingo are increasingly advocating users to have tests and retrieval, rather than restudy. Previous work has attributed the testing effect to myriad factors, such as retrieval effort (6, 7) and elaborative semantic organization during testing (8, 9), which are absent during studying. Nevertheless, its underlying mechanisms remain a topic of debate. Here, we report and test an emergent predictive learning account as its cognitive and neural origin (10–12).

Predictive learning implies that minimizing prediction errors is a key objective for learning (13). This is a foundational principle in computational approaches to learning (including in Artificial Intelligence) and has more recently found its way into human declarative memory as well. Indeed, several studies have demonstrated that prediction errors can significantly promote declarative memory (14–16). For example, a student might initially predict that a dolphin is a type of fish, but is then corrected that it is, in fact, a mammal. Here, prediction errors can restructure and cement the student’s information (neural representations) in memory.

Of importance for the present purpose, prediction error is more likely to appear in testing, relative to studying (10). Indeed, testing provides more opportunities to predict possible answers, relative to studying. For example, students have no opportunity to guess (predict) which city is the capital of Australia when the teacher provides the answer directly. The mismatch between predictions and subsequent, external or internal, feedback generates prediction errors driving learning in a testing context (10, 12). External feedback comes from the environment, while internal feedback can manifest as internal (confidence) feelings about whether a prediction is correct (17, 18).

Indeed, the neural signature of external feedback can also be active when people have an internal feeling of being correct or incorrect (18–20). Importantly, both external and internal prediction errors are consistently localized in the midbrain across animal species including rodents (21), macaques (22, 23), and humans (17, 24), where they are encoded by dopaminergic bursts. Recent studies suggest that the effect of prediction error on declarative memory can be fully mediated by the neural activation in the ventral striatum (25), a key target for dopamine release. Based on these results, we propose a dopaminergic neural basis (ventral striatum) for the testing effect and suggest that this neural basis can be explained from a predictive learning perspective. To substantiate this account, we employed a combination of cognitive neuroscience (fMRI) experiments and computational modeling techniques to investigate whether both testing and prediction errors elicit ventral striatum activation, and whether such activation enhance declarative memory.

## Results

### Experimental design

Our declarative memory task consisted of four phases designed to help participants learn 90 Dutch-Swahili word pairs (Figure 1A) (10). In Phase 1, participants underwent initial learning, during which each word pair was displayed on the screen for 3 seconds. This phase ensured that participants acquired initial knowledge for the subsequent tests. Phase 2 involved a no-feedback assessment to control participants’ initial learning performance. During this phase, participants selected the correct Swahili translation for a given Dutch word from four options and rated their confidence in their choice. A previous study used strategies such as excluding correct or high-confidence items from Phase 2 to filter out extreme initial learning (10). However, here and in previous study (10), the behavioral patterns remain unchanged with or without applying these filters (See Supplementary Figure S1). For this reason, and to maintain strong statistical power in our fMRI analyses, we chose not to filter. The primary variable, Test vs. Study, was manipulated in Phase 3. Participants selected the correct Swahili translation of a Dutch word, either from four boxed options (test trials) or from a single boxed option containing the correct answer (study trials). After making their selections, participants rated their confidence and received (informative but non-monetary) feedback with the correct answer. Importantly, participants were informed in advance that the solely boxed option in each study trial was always correct, and they were only allowed to select this correct option. Therefore, they were always confident in their choices and experienced no (or the smallest possible) prediction errors in the studying condition. Phase 4 consisted of two final assessments without feedback. A word pair was considered fully learned only if it was correctly identified in both final assessments*. Only Phase 3 was conducted in the fMRI scanner.

**Figure 1.**
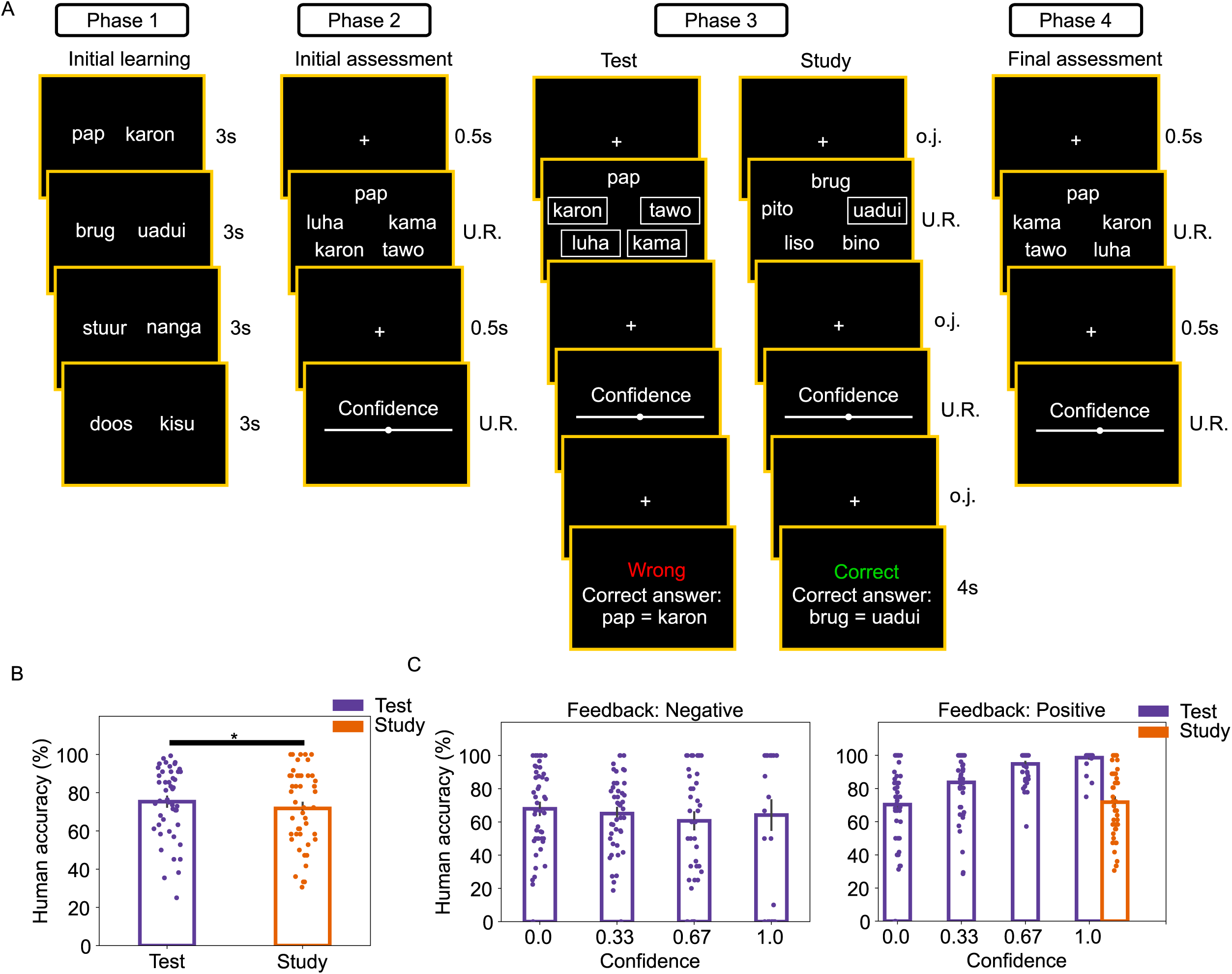
Experimental procedure (A) and behavioral results (B-C). **A.** Participants completed a four-phase task to learn 90 Dutch-Swahili word pairs. Phase 1: All word pairs were presented randomly, with each pair displayed for 3 seconds to facilitate initial learning. Phase 2: An assessment without feedback was administered to assess initial learning. In each trial, participants selected the Swahili translation of a Dutch cue from four options and rated their confidence in being correct (Confidence scale simplified for illustration purposes). Phase 3: This phase introduced the Test vs. Study manipulation and prediction errors. In testing trials, participants selected the Swahili translation from four boxed options, while in studying trials, they could only choose the boxed correct answer. After each choice, they rated their confidence and received feedback, including whether they were correct and the correct answer. Importantly, three optimized jitter intervals were inserted between the choice onsets, confidence rating, and feedback to ensure precise timing and minimize expectancy effects. Phase 4: Two final assessments, using the same procedure as Phase 2 (test without feedback), were conducted to evaluate final learning performance. **B.** Behavioral pattern in the current study replicated the robust testing effect. **C-D.** The testing effect was still robust after controlling feedback valance and confidence ratings from Phase 3. Individual data points are represented by the dots; The error bars reflect the standard errors; Confidence ratings were from Phase 3; *p* < 0.05*. *N* (subjects) = 48.

### Robust testing effect in behavioral data

As shown in Figure 1B, the final assessment accuracy (Phase 4) after test is better than that after study, which replicates the robust testing effect (*χ^2^*(1, N = 48) = 6.382, *p* =.012, β *± 95% CI* =.036 ±.029, *R^2^* =.001). The testing effect became more robust (*χ^2^*(1, N = 48) = 81.702, *p* <.001, β *± 95% CI* =.225 ±.050, *R^2^* =.078) after controlling for feedback valence and confidence ratings from Phase 3 (Figure 1C).

### Predictive learning model accounts for the behavioral data

We next turned to computational modeling and constructed an associative memory neural network model (Figure 2A; see Methods) where Dutch and Swahili words were presented as input and output, respectively (10). Initial connections between Dutch and Swahili units were based on participants’ confidence ratings from Phases 2 and 3. The model then underwent testing and studying trials based on predictive learning, Hebbian learning, or a combination of both. Hebbian learning learns from feedback. We included Hebbian learning as a baseline model to evaluate empirical fit of a model that does not learn from prediction error. Finally, the model completed a simulated final assessment to evaluate its performance. Based on different combinations of learning components (initial learning, predictive learning, and Hebbian learning), a total of seven models were constructed (Table 1).

**Figure 2.**
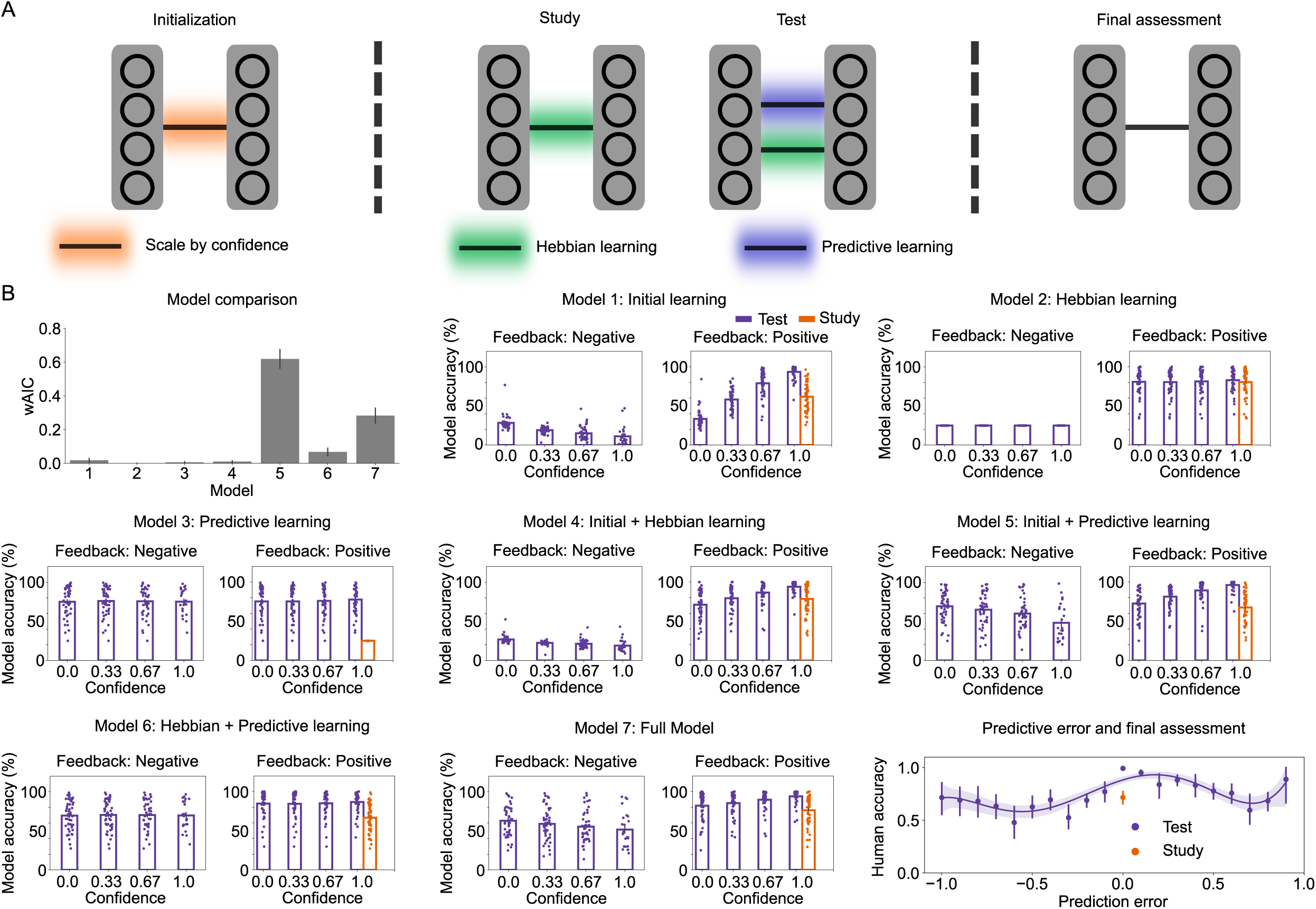
Model architecture (A) and modeling results (B). **A.** An associative memory neural network with 90 Dutch input units and 360 Swahili output units was developed to simulate human learning behavior. The model’s initial connections were scaled according to human confidence ratings from Phase 2 and Phase 3. After initial connections, the model went through testing and studying trials as in Phase 3 of the human task. The model updated connections via Hebbian learning during both studying and testing trials. Predictive learning, however, was applied exclusively during testing trials. Final assessment was conducted as in Phase 4 of the human task. **B.** The model comparison with wAIC (higher means better) and model simulations suggested that the model (Model 5) with both initial learning and predictive learning was the best model. Moreover, only models with predictive learning (Models 3, 5, 6, 7) could replicate the testing effect. Bottom right panel: Both extremely positive (around 1) and negative prediction errors (around-1) enhance the final assessment accuracy. The peak point around small positive prediction errors (around 0.1) was triggered by good initial learning. Individual data points are represented by the dots; The error bars reflect the standard errors; Confidence ratings were from Phase 3. *N* (subjects) = 48.

**Table 1:**
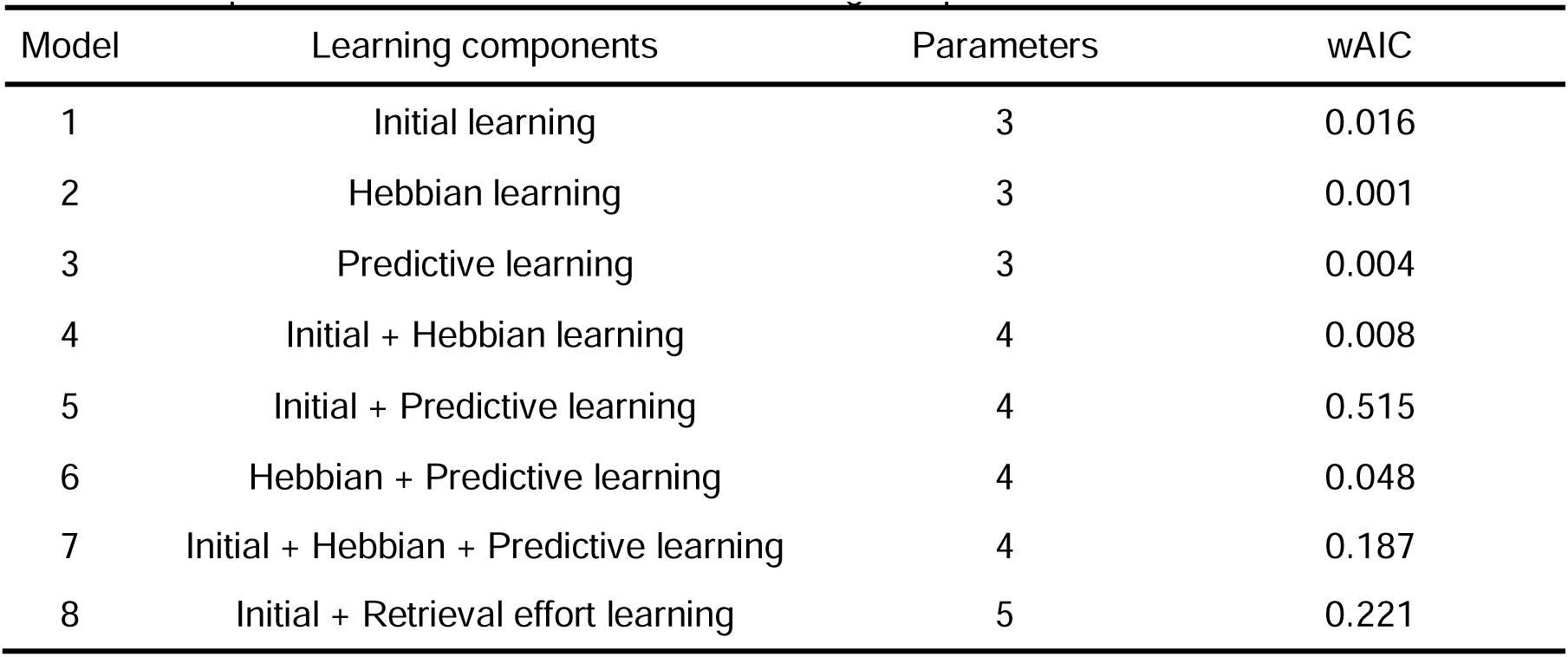
Computational models with different learning components.

Model comparison with wAIC (higher means better) suggested that the model with both initial learning and predictive learning (Model 5) fit the human data best (Figure 2B). Replicating previous research (10), model simulations showed that only the models incorporating predictive learning (Models 3, 5, 6, 7; Figure 2B) could replicate the testing effect in human data.

Additionally, we developed an enhanced Hebbian learning model that incorporates an implementation of the retrieval effort hypothesis (6, 7). That is, greater learning occurs following difficult but successful tests, compared to easier successful tests or study. In our experiment, confidence ratings serve as a proxy for difficulty. Accordingly, we specified a minimal baseline learning rate applied across all trials, with the learning rate in each trial inversely modulated by correct confidence ratings (i.e., higher learning rates for low-confidence correct trials; see Methods). Under this scheme, both incorrect testing trials (with wrong confidence ratings) and studying trials (with maximum confidence ratings) were assigned only the minimal learning rate. However, model comparisons still favor a model that combines initial learning and predictive learning (Model 5; Supplementary Figure S2, Panel A). Moreover, the model combining initial learning and retrieval effort (Hebbian) learning did not show a clear distinction between testing and studying conditions (Model 8; Supplementary Figure S2, Panel B).

Finally, we estimated trial-level prediction errors for each participant from the best-fitting model (Model 5) and examined their relationship with final assessment accuracy (bottom right panel in Figure 2B). For clarity in visualization, prediction error values were rounded to one decimal place. The influence of prediction errors on final assessment accuracy followed a W-shaped pattern, highlighting three key peaks. The first peak occurred at extremely negative prediction errors (around-1), and the second at extremely positive prediction errors (around 1), consistent with a critical role of (unsigned) prediction errors in memory consolidation (15, 26, 27). The third peak, at small positive prediction errors (around 0), was driven by initial learning rather than prediction errors. Specifically, strong initial learning enabled participants (and the model) to consistently provide correct answers in both Phase 3 and Phase 4, resulting in small prediction errors in Phase 3 but high final assessment accuracy in Phase 4. Indeed, the initial learning effect is obvious in the human behavioral pattern, in which participants’ final assessment accuracy increases with correct confidence ratings in Phase 3 (Figure 1C). Importantly, this W-shaped relationship was consistent across all relevant datasets, including the behavioral data from the current fMRI study and two behavioral datasets with the same experimental design from a previous study (10) that we re-analyzed (see Supplementary Figure S2, Panels C-D).

### Testing triggers neural activation in the ventral striatum, insula, and midbrain

The fMRI analysis first examined neural activation associated with testing (relative to studying) during the feedback onsets of Phase 3. In the Test-GLM1 (see Methods), we included three main regressors (correct test, incorrect test, and study) separately for the feedback onsets and choice onsets (a total of 6 main regressors). Consistent with our assumption, testing elicited stronger activation than studying in the ventral striatum (VS; Figure 3A). Additionally, the insula, midbrain, superior and middle frontal gyrus (SFG and MFG) also showed greater activation during feedback onsets following test, compared to study (Figure 3A). Notably, both correct (Figure 3B) and incorrect (Figure 3C) test trials showed greater activation in the VS, insula, and midbrain compared to study trials, indicating that these brain responses are not merely driven by positive feedback in correct tests (or negative feedback in incorrect tests). In the following fMRI analyses, we mainly focus on VS, insula, and midbrain, as other testing-related regions such as MFG and SFG did not consistently emerge in both correct and incorrect tests.

**Figure 3.**
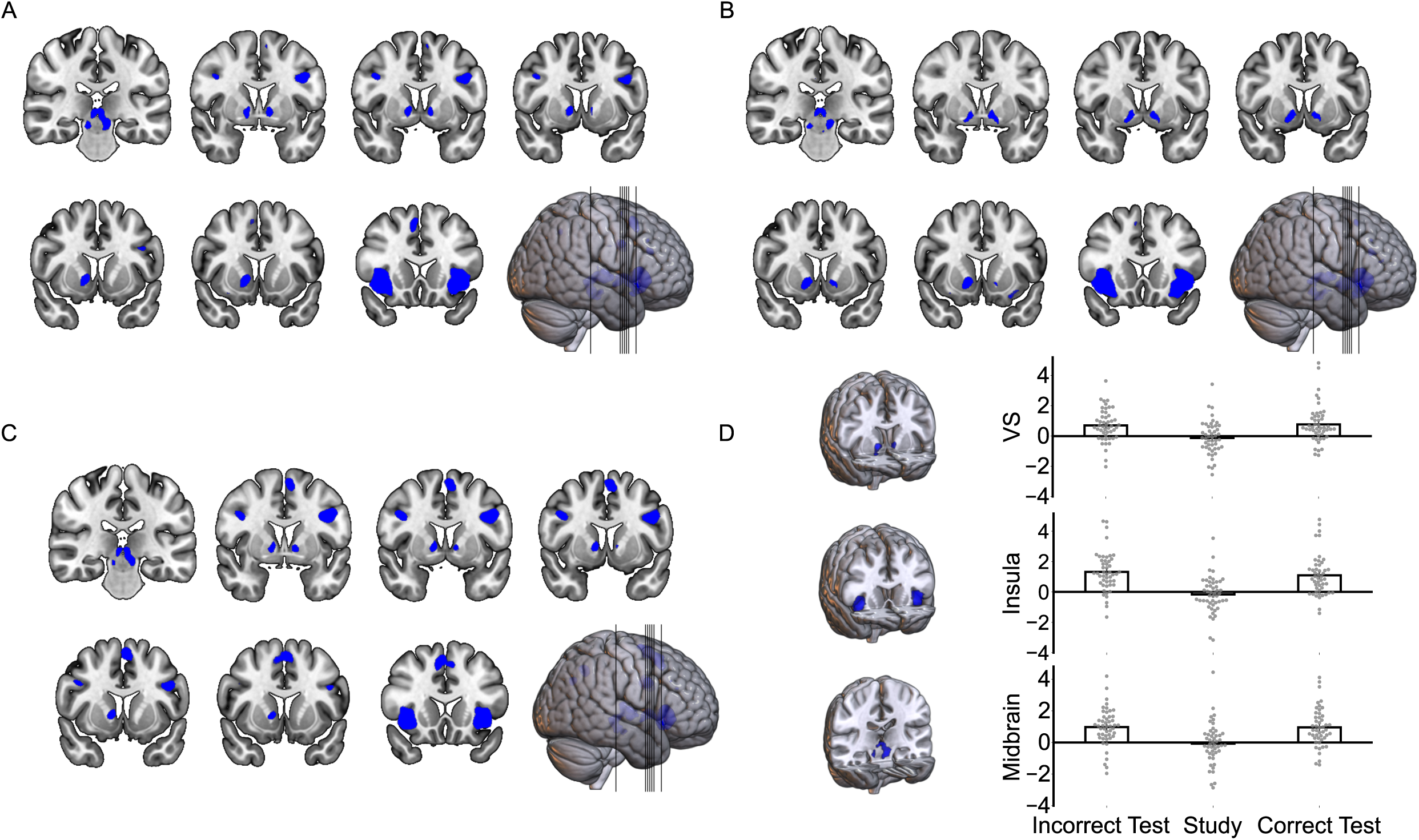
Test (vs Study) related neural activations during feedback onsets. A-C: Test (vs Study; A), Correct test (vs Study; B), and Incorrect Test (vs Study; C) all revealed increased neural activation in the VS, insula, and midbrain during feedback onsets. **D**. The test-related activation in the VS, insula, and midbrain were defined as regions of interest (ROIs). The mean activations in these ROIs were high for both correct and incorrect test trials, but low for study trials. Individual data points are represented by the dots; The error bars reflect the standard errors. *N* (subjects) = 48.

As an exploratory analysis, we also examined neural activity during choice onsets, where testing (relative to studying) elicited significant activation in the VS and visual cortex (Supplementary Figure S3, Panel A). The visual cortex activation may reflect visual search (i.e., selecting the correct answer among four options), while the VS activation may reflect active anticipation.

Consistently, in another Test-GLM2 (see Methods) that included trial-level confidence ratings as a parametric modulator for the (testing) choice onset regressors, we found that confidence positively modulated VS activation (Supplementary Figure S3, Panel B). This finding aligns with prior evidence that dopaminergic activity increases when agents anticipate a positive outcome (22, 23), and with a previous study showing that the VS is also active when participants feel they have grasped the meaning of a new word (28).

To ensure that the VS activation observed during feedback onsets was not simply residual VS activation from the preceding choice onsets, we examined the testing (vs. studying) related neural activity during two fixation periods between choice and feedback onsets (Additional regressors in Test-GLM1; see Methods). However, no fixation periods showed increased VS activation following test compared to study (Supplementary Figure S3, Panels C-D).

#### Prediction error also triggers neural activation in the VS, insula, and midbrain

Our theory also predicts an overlap between prediction error and testing-related signals in the VS. Accordingly, in the PE-GLM1, trial-level prediction errors and feedback valence were jointly included as parametric modulators of one feedback onset regressor (See Methods). Neural activity positively modulated by prediction error (Figure 4A) was localized in the VS, insula, and midbrain. In contrast, the feedback valence parametric modulator revealed non-overlapping neural signatures in the anterior cingulate cortex (ACC) and posterior cingulate cortex (PCC; Figure 4B). We conclude that the VS, insula, and midbrain are sensitive to prediction errors, rather than to feedback valence. Importantly, the neural activation elicited by testing and prediction error overlap in the VS, insula, and midbrain (Figure 4C).

**Figure 4.**
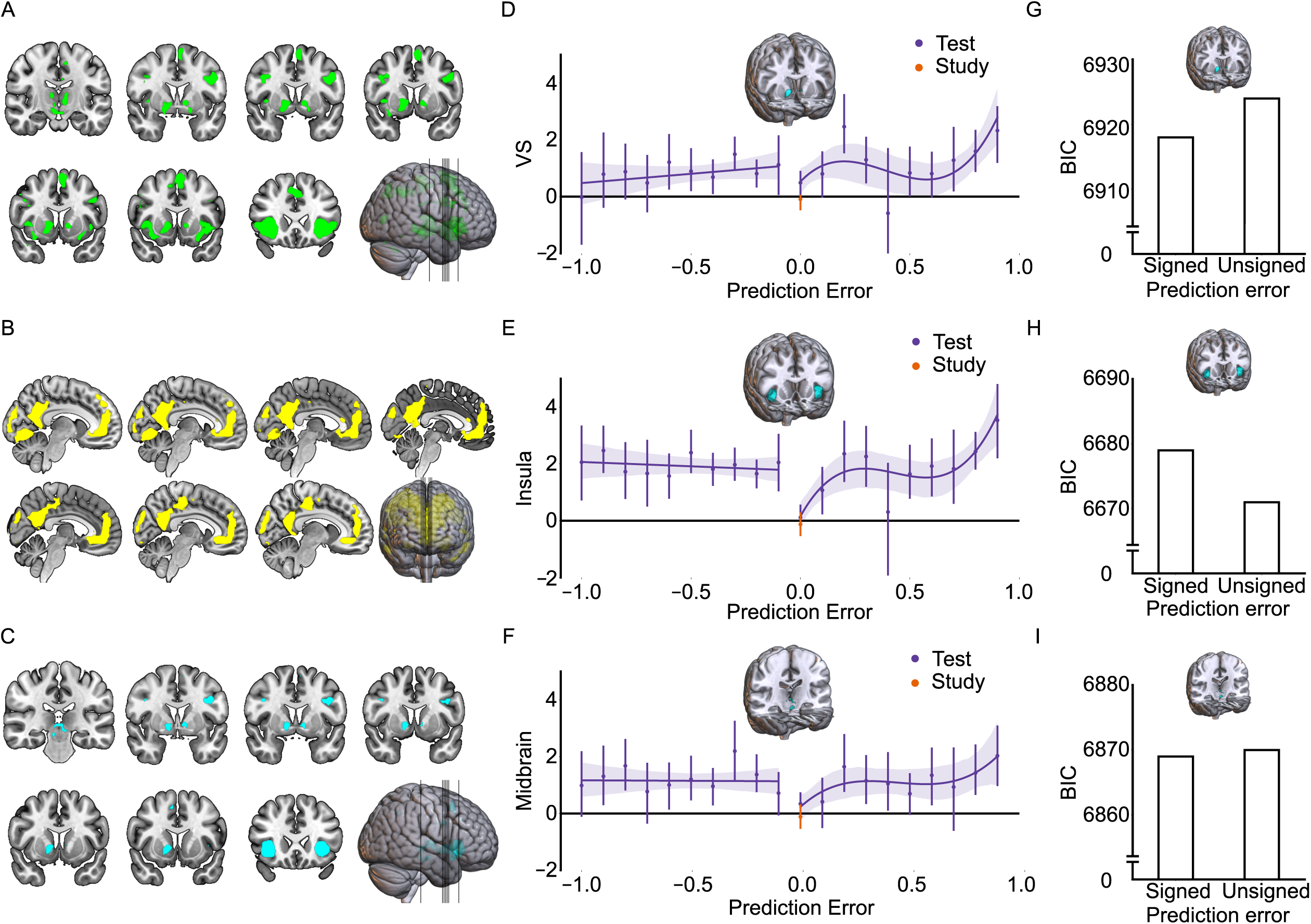
Neural signatures of prediction errors. A: The VS, insula, and midbrain activations increase with the prediction errors during the feedback onsets. **B:** Positive feedback elicited stronger neural activations in ACC and PCC, relative to negative feedback. **C:** Testing and prediction errors elicited overlapping neural activations in the VS, insula, and midbrain. **D-F:** The relationship between prediction errors and mean neural activation (across participants) in the VS, insula, and midbrain ROIs (overlapping regions between testing and prediction errors). VS activation tracked signed prediction errors (Panel D), insula activation reflected unsigned prediction errors (Panel E), and midbrain activation showed a better fit with signed prediction errors (Panel F). The error bars reflect the standard errors. **G-I:** Signed or unsigned prediction errors were included into separate general linear mixed models to predict neural activity within the VS, insula, and midbrain regions of interest (ROIs). Signed prediction errors better predicted activation in the VS (Panel G; lower BIC means a better fit) and midbrain (Panel I), whereas unsigned prediction errors better predicted activation in the insula (Panel H). Because the BIC values are large, the y-axes in Panels G-I are truncated to improve visibility. Axis breaks (=) are used to indicate the truncation. *N* (subjects) = 48.

To examine how the VS, insula, and midbrain encode prediction errors, we defined regions of interest (ROIs) based on overlapping VS, insula, and midbrain regions between prediction error and testing. We then constructed another PE-GLM2, in which feedback onsets associated with different prediction errors (rounded to one decimal place for statistical robustness of the estimates) were modeled as separate regressors (See Methods). For each participant, we extracted the mean beta values across voxels within the VS, insula, and midbrain ROIs for each regressor. Interestingly, VS activity seems to track signed prediction errors (Figure 4D), whereas insula activity reflects the absolute (unsigned) magnitude of prediction errors (Figure 4E). Indeed, when using either signed or unsigned prediction errors in separate general linear mixed models (separate signed model and unsigned model; with the PE-GLM2 betas as dependent variables; the feedback valence was always included as a covariate) to predict neural activity within the VS, insula, and midbrain ROIs, we observed different patterns. VS activation was primarily predicted by the signed prediction errors (Figure 4G; Lower BIC indicates better model fit; Signed prediction error: *χ^2^*(1, N = 48) = 10.414, *p* =.001, β *± 95% CI* =.146 ±.089, *R^2^* =.007; Unsigned prediction error: *χ^2^*(1, N = 48) = 4.605, *p* =.032, β *± 95% CI* =.060 ±.055, *R^2^* =.003). In contrast, insula activation was better explained by unsigned prediction errors (Figure 4H; Signed prediction error: *χ^2^*(1, N = 48) = 31.561, *p* <.001, β *± 95% CI* =.247 ±.086, *R^2^* =.034; Unsigned prediction error: *χ^2^*(1, N = 48) = 39.674, *p* <.001, β *± 95% CI* =.171 ±.053, *R^2^* =.039). Additionally, midbrain activity was slightly better predicted by signed prediction errors (Figure 4I; Signed prediction error: *χ^2^*(1, N = 48) = 8.464, *p* =.004, β *± 95% CI* =.132 ±.089, *R^2^* =.010; Unsigned prediction error: *χ^2^*(1, N = 48) = 7.589, *p* =.006, β *± 95% CI* =.078 ±.056, *R^2^* =.009).

Notably, neither VS nor insula encoded positive prediction errors in a strictly linear fashion. When the prediction error was close to zero, both regions exhibited minimal activation. Both VS and insula activations peaked around a prediction error of 0.2, possibly reflecting a state in which participants strongly anticipated the positive outcome and received confirmatory feedback. A second peak in VS and insula activation was observed at the large prediction error range (around 1), reflecting neural responses to highly unexpected positive feedback.

### VS, insula, and midbrain influence the final assessment accuracy

Finally, we conducted a back sorting analysis (Bsort-GLM1) by categorizing feedback onsets in Phase 3 based on final assessment accuracy in Phase 4 (feedback onsets followed by correct vs. incorrect Phase 4 assessments), modeling them as two main regressors in the Bsort-GLM1 (see Methods). To minimize the confounding effects of initial learning on final assessment, we included only incorrect trials and correct trials associated with high prediction errors (prediction error ≥ 0.7) from Phase 3, as these trials were less likely to be influenced by the initial learning. Consistent with our assumption, the VS (and dorsal striatum), insula, and midbrain showed increased activation during feedback onset when the following final assessment was correct (relative to incorrect; Figure 5A), overlapping with VS and insula regions elicited by testing and prediction error (Figure 5B-D).

**Figure 5:**
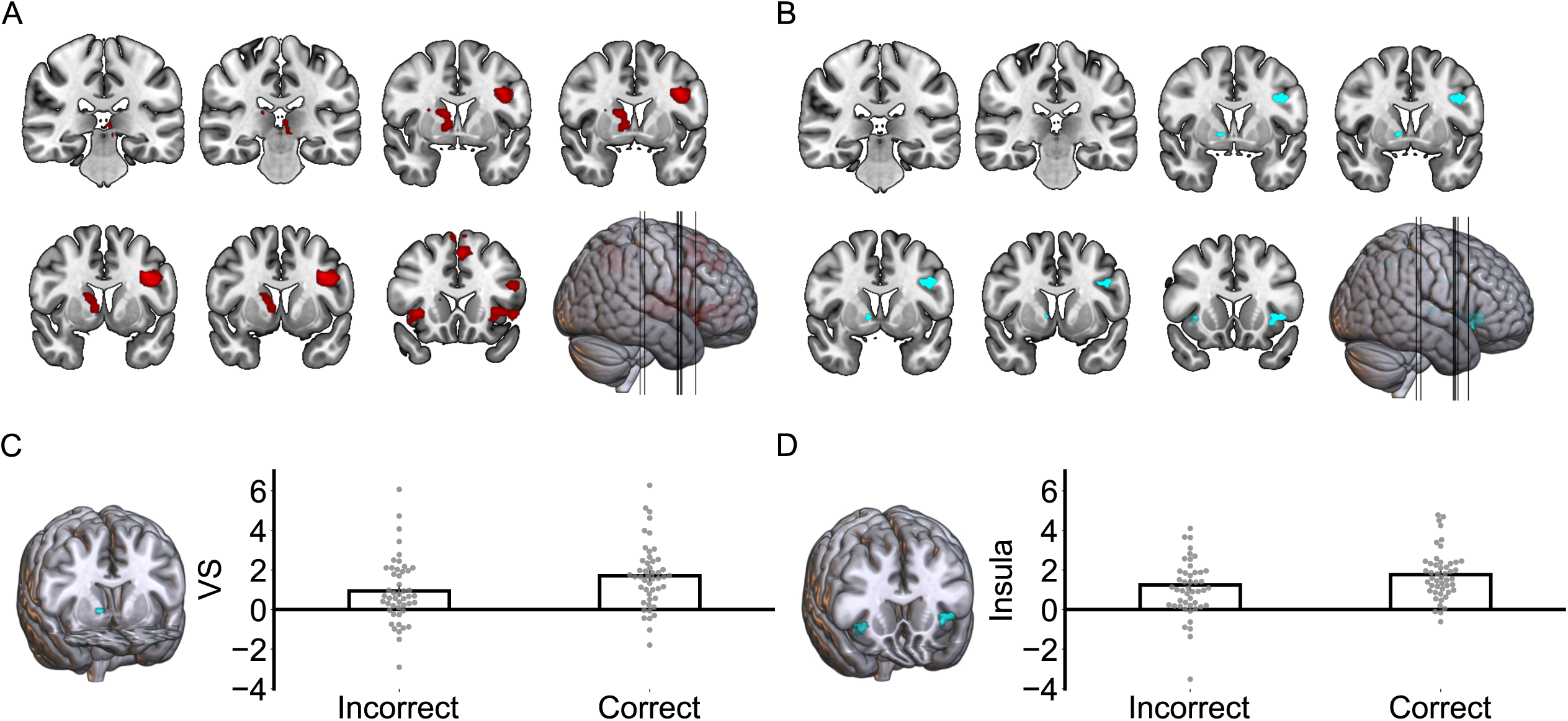
N**e**ural **signatures of final assessment accuracy during feedback onsets. A:** Activation in VS (and dorsal striatum), insula, and midbrain during feedback onsets was associated with final assessment accuracy. **B:** Neural activation related to testing, prediction errors, and final assessment accuracy overlapped in the VS and insula. **C-D:** The overlapping regions in the VS and insula were defined as ROIs. Mean activation within these ROIs were higher during feedback onsets when the subsequent final assessment was correct, compared to when it was incorrect. Individual data points are represented by the dots; The error bars reflect the standard errors. *N* (subjects) = 48.

To ensure the robustness of our findings, we conducted an additional Bsort-GLM2 that included again the incorrect trials and correct trials (with a looser threshold of prediction error ≥ 0.5) from Phase 3. This analysis still revealed VS and insula activations that overlapped with those associated with testing and prediction error (Supplementary Figure S3, Panels E-F). Moreover, another exploratory Bsort-GLM3 targeted at choice onsets revealed no stronger brain activation for correct compared to incorrect final assessment. The latter result demonstrates that the effect observed in VS and insula is, as predicted, specific for the feedback onsets.

## Discussion

This study combined behavioral, modeling, and fMRI approaches to explore predictive learning as a potential account for the testing effect. The behavioral and modeling results reveal (unsigned) prediction errors as a key driver of the testing effect. Further fMRI results suggest that both Test (vs Study) and prediction error activate VS, insula, and midbrain during feedback onsets.

Importantly, the VS encoded signed prediction errors, while insula encoded unsigned prediction errors. Furthermore, activations in the VS and insula were associated with the final assessment accuracy.

### Predictive learning as a behavioral explanation of the testing effect

Our computational modeling results indicate that the testing effect in human data could only be explained by an associative memory neural network model that incorporates predictive learning, but not Hebbian learning. This aligns with the computational principle that predictive learning is more efficient than Hebbian learning (29). Unlike Hebbian learning, predictive learning requires active retrieval or prediction, consistent with the cognitive and educational trends that advocate for active, prediction error-based learning (30, 31). Augmenting the Hebbian learning model with retrieval effort modulation (6, 7) did not enhance its explanatory power beyond that of the predictive learning model. However, it is worth noting that the retrieval effort theory (6, 7), along with several other previous explanations (extra semantic organization or extra feedback during test compared to study) of the testing effect (5, 8, 32), share key assumptions with our predictive learning account, so they may eventually be integrated within a broader predictive learning framework. First, consistent with the retrieval effort assumption, individuals may experience strong prediction errors during tests that are difficult but ultimately successful (6, 7). Second, predictive learning may help organize an elaborative semantic network (29, 33), a possibility that future studies could further investigate. Finally, the exact function of feedback is to trigger prediction errors in a testing context.

The trial-level prediction errors estimated by the optimal predictive learning model influences final assessment accuracy in a W-shaped pattern. This pattern suggests an unsigned prediction error effect that both extremely positive and negative prediction errors enhance declarative memory (15, 34). Notably, extremely negative prediction errors are triggered by high-confidence errors, which is labeled the hypercorrection effect (26, 35–37). The hypercorrection effect suggests that high-confidence errors are more easily corrected than low-confidence errors (37). Although several studies support the hypercorrection effect (26, 35, 36), a recent study has reported contradictory findings, indicating that small errors are easier to correct (38). Considering our findings, one possible explanation is that the study failing to observe the hypercorrection effect might not elicit extremely negative prediction errors.

It is important to clarify that while prediction errors are more likely to occur in the testing condition, they are not exclusive to testing and may also arise during study. For example, if people distrust the provided answer or unexpectedly receive negative feedback during a studying trial, they may also experience high prediction errors and potentially enhanced declarative memory. We did not include such manipulations in our study to avoid blurring the distinction between testing and studying conditions. However, we believe that predictive learning may serve as a general learning rule across different contexts (both testing and studying), as long as active prediction is encouraged. In other words, the most effective learning strategy may not necessarily be testing, but rather continuously engaging in critical thinking and active prediction, even when the “correct answer” has already been revealed.

### Testing effect and prediction error effect share the same neural basis

Our fMRI results revealed that both test and prediction errors robustly activate VS, insula, and midbrain during feedback onsets, which reveals a shared dopaminergic basis between testing and prediction errors. Indeed, the VS is a key target of dopaminergic transmission (39), and the midbrain serves as a primary source of dopamine release (21, 40). Dopaminergic signals are broadly transmitted across brain regions, including VS, hippocampus, and neocortex (39, 41), forming a distributed learning system (25, 28). Previous studies have reported VS activation and its functional connectivity with neocortex when participants successfully acquire the meaning of a novel word (28). Additionally, VS-hippocampus connectivity emerges during Swahili word learning and is modulated by reward prediction errors (25), suggesting that prediction errors play a crucial role in activating the above complementary learning system. Future studies can explore how this complementary dopaminergic learning system promotes testing and prediction error-based learning.

In our original theoretical framework, we largely overlooked the important role of insula. The insula also has intensive dopaminergic receptors (42–44). Besides, a recent study suggests that the insula serves as a top-down modulator of the dopaminergic release in the midbrain (45).

Importantly, the insula receives not only dopaminergic afferents, but also cholinergic, serotonergic, and noradrenergic inputs (44), suggesting a potential interplay among multiple neuro-modulatory systems.

Consistent with previous findings, VS and insula encode prediction errors in distinct ways (25, 46). Specifically, VS activation encodes the original prediction errors as signed prediction errors (25), while the insula encodes the absolute magnitude of the prediction errors as unsigned prediction errors (46). Given that final assessment accuracy was influenced by unsigned prediction errors, the insula may serve as an important neural hub in the current study.

Interestingly, both VS and insula exhibited two peak points in response to positive prediction errors. The first peak point occurs at a low but non-zero prediction error range, aligning with a recent finding that participants are most curious when they are close to the correct answer (47). The second peak point emerges in the large prediction error range, likely reflecting surprise to unexpected positive feedback. Future studies could investigate how these peak points influence learning in different tasks, offering important insights for both cognitive research and educational practice. Future modeling studies could explore whether incorporating complementary systems for signed and unsigned prediction errors, along with dynamic learning rates that peak at the above points, can enhance learning efficiency.

Our study offers a piece of puzzle into the ongoing debate over whether declarative memory is influenced by signed versus unsigned prediction errors (48). First, our behavioral results, together with previous studies on the hypercorrection effect (36, 37), suggest that internally generated, extremely negative prediction errors may be critical for eliciting unsigned prediction error effects. Indeed, prior studies demonstrating signed prediction error effects typically manipulated prediction errors through external cues, rather than through participants’ internal confidence (14, 49, 50). Second, our neural findings suggest a potential dissociation in prediction error encoding between the VS and insula. Complementing this spatial dissociation, a previous EEG study reported that signed and unsigned prediction errors are encoded by late and early EEG components, respectively, during the feedback stage (51). Taken together, these findings indicate that signed and unsigned prediction errors may coexist within spatially and temporally dissociable neural systems.

### Internal feedback as a potential explanation of the testing effect

In our study, prediction error was elicited by external feedback. However, the testing effect can occur in the absence of external feedback, leading to a classification of testing effects with external feedback (52, 53) and those without (3, 7). Although the current predictive learning framework can account for the testing effect with external feedback, it requires modifications to explain the testing effect without external feedback. As mentioned earlier, some neural studies have shown that internal feedback (e.g., confidence) can also elicit VS activation and modulate learning (18, 28). Moreover, post-test confidence rating can replace reward signal to elicit VS-encoded internal prediction errors (17). Consistent with these studies, our study also found that confidence modulated the VS activity during testing. Therefore, predictive learning may contribute to the testing effect by providing both internal and external feedback and prediction errors.

However, because initial learning is inherently confounded with confidence, it remains challenging to disentangle the effect of initial learning from learning driven by internal feedback (confidence), which may be an important topic for future research.

## Summary

In summary, the current study supports the notion that the testing effect benefits from predictive learning, as evidenced by cognitive neuroscience and modeling findings. Notably, the testing effect represents just one instance of a broader range of active learning strategies that emphasize active predictions (54, 55). Predictive learning may not only serve as a specific mechanism underlying the testing effect but also offer a broader cognitive framework for general active learning approaches, such as the generation effect (54), problem-based learning (30, 31), and error-driven learning (26, 55).

## Materials and Methods

### Participants

Fifty-five participants, all native Dutch speakers, with no reports of smoking, alcohol consumption, medication use within 24 hours of scanning, or psychiatric disorders, were recruited for this study. Each participant was compensated 35€. Data from seven participants were excluded from the final analysis: six due to technical issues during the experiment, and one due to being left-handed. This resulted in a final sample of 48 right-handed participants (mean age = 22.44, SD = 2.65, range = 18-29; 34 female participants, 14 male participants). The study was approved by the local ethics committee at Ghent University Hospital, and all participants provided informed consent.

### Experimental design

Participants were instructed to learn 90 Dutch-Swahili word pairs (Figure 1A). They underwent 4 phases: Initial learning (Phase 1), Initial assessment (Phase 2), Test vs Study (Phase 3), and Final assessment (Phase 4). Given our primary interest in the neural mechanism underlying Test vs Study comparison, only Phase 3 was conducted in the fMRI scanner.

#### Phase 1: Initial learning

Participants were required to familiarize themselves with all word pairs before proceeding to the subsequent phases. All word pairs were randomly presented on the screen, with each pair displayed for 3 seconds.

#### Phase 2: Initial assessment

Following initial learning, a test without feedback was conducted in Phase 2 to assess participants’ learning progress before the formal manipulations in Phase 3. In each trial, participants selected the Swahili translation of a randomly presented Dutch word from four random options by pressing “f”, “v”, “n”, or “j” on the keyboard, followed by a 4-point confidence rating (“very uncertain,” “uncertain,” “certain,” or “very certain”) for their choice.

#### Phase 3: Test vs Study

Phase 3 was the critical phase, designed to manipulate the Test vs Study conditions and induce prediction errors. In each trial, participants were presented with a randomly selected Dutch cue and four random Swahili options, some of which were boxed. Participants could only choose from the boxed options, provide a confidence rating, and then receive feedback indicating whether their choice was correct (“Correct” in green or “Wrong” in red), along with the correct answer.

Responses were made using a Cedrus Lumina LS-Pair MRI-compatible ergonomic response pad, where participants used their left middle finger, left index finger, right index finger, and right middle finger to select corresponding options. In testing trials, participants selected from four boxed Swahili options, allowing them to predict the possible answer and experience the prediction error when feedback was provided. In contrast, during studying trials, the correct Swahili option was the only one boxed, leaving no opportunity for prediction errors as participants could only select the correct answer. Among the 90 trials in Phase 3, 72 trials (80%) were assigned to the testing condition, and 18 trials (20%) to the studying condition. We included more testing trials because the testing condition involves correct and incorrect trials (and different prediction errors), which requires enough trials for reliable analysis. The exact number of correct and incorrect testing trials varied across participants, depending on initial learning. After data collection, each participant, on average, had 37.75 correct (52.43%) and 34.25 incorrect testing trials (47.57%) in Phase 3, which was nearly balanced. All word pairs were randomly assigned to testing and studying conditions, and the trial order was randomized for each participant. After data collection, the mean confidence rating (in Phase 2) was 0.388 in the testing condition and 0.389 in the studying condition, indicating that initial learning was well counterbalanced between the two conditions. Additionally, three fixation intervals with optimized jittering were inserted before the onsets of choices (Jitter 1), confidence rating (Jitter 2), and feedback (Jitter 3). Since feedback onset was the most critical, Jitters 1 and 3 were longer than Jitter 2. Specifically, Jitters 1 and 3 were sampled from a Poisson distribution with a mean of 4.35 seconds (ranging from 3 to 8 seconds), while Jitter 2 was sampled from a shorter Poisson distribution with a mean of 2.35 seconds (ranging from 1 to 6 seconds).

#### Phase 4: Final assessment

Two final assessments were conducted to evaluate the learning performance, following the same procedure as in Phase 2. Administering two assessments was diagnostic to know whether participants selected the correct answer based on actual knowledge rather than chance.

Specifically, a Swahili answer was considered fully learned only if a participant selected this correct answer in both assessments, which is unlikely to occur by chance on two consecutive assessments.

### Behavioral data analysis

A Generalized Linear Mixed Model (GLMM) was employed to analyze the trial-level data. The predictors in the model included Test vs. Study (Test = 1, Study = 0), Feedback (Positive = 1, Negative = 0), and confidence ratings (0, 0.33, 0.67, 1). The model incorporated subject-specific random slopes for predictors and a random intercept^†^. Final assessment accuracy was the dependent variable, coded as 1 for correct responses in both final assessments (Phase 4 and Phase 5), 0.5 for one correct response^‡^, and 0 for no correct response^§^.

### Modelling analysis

#### Model architecture

Several neural networks incorporating Hebbian learning and/or predictive learning were developed to fit the behavioral data (Figure 2A). By adjusting the combination of learning components within the model, we assessed which components were critical for replicating the testing effect. For instance, if the testing effect could only be replicated when predictive learning was included, it suggests that predictive learning is essential for the testing effect. Besides, if the predictive learning model is the best-fitting model, its optimized parameters would be used to estimate trial-level prediction errors in human data, which would then be applied in the fMRI analysis to investigate the neural basis of prediction errors.

The model’s task was to establish accurate connections between 90 Dutch input units and 360 Swahili output units. Its initial connections were aligned with participants’ confidence ratings from Phase 2 and Phase 3, simulating the initial learning from Phase 1 (10). The exact equation is shown below:

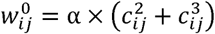

In this equation, cij^2^ and cij^3^ represent the confidence ratings from Phase 2 and 3, respectively.

The relationship between confidence ratings and initial connections was scaled by parameter α. In the study condition, participants were given the correct answer directly and could not express their internal confidence in Phase 3, meaning we had no confidence data for those items in Phase 3. To address this, we substituted study-trial confidence *c_s_*with the mean confidence rating from the testing condition:

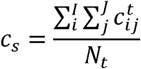

for each studied (rather than tested) word.

After establishing the initial connections, the model proceeded with testing and studying trials like Phase 3 in human task. In testing trials, the model selected the correct Swahili translation of a Dutch word from four options, while in the studying trials, the model could only choose the correct Swahili answer. The connections between Dutch and Swahili words were updated using Hebbian learning or/and predictive learning. Hebbian learning strengthened the Dutch and Swahili target connections in reward contexts (positive feedback trials), while predictive learning adjusted the connections based on prediction errors.

Regardless of which learning component was applied, the model generated its predictions according to the following equation:

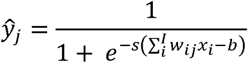

Thus, the activation of each element 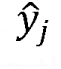 in the output layer was determined by the weighted (w_ij_) sum of the input values x_i_, which was then passed through the sigmoid function. Non-option output units always have zero activations in a trial.

Hebbian learning, modulated by reward, was implemented as follows:

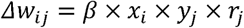

In this equation, the model updated the weight w_ij_ only when a reward r_j_ was present (i.e., when feedback was positive). Note that Hebbian learning not modulated by reward would promote learning in testing and studying conditions equally, thus was impossible to explain the testing effect.

Hebbian learning incorporated retrieval effort assumption (7) was implemented as follows:

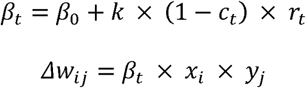

Here, β_O_ represents the minimal learning rate applied in each trial. Building on this baseline, the exact learning rates were inversely modulated by correct confidence ratings (scaled by *k*).

Consequently, incorrect testing trials (with wrong confidence ratings) and studying trials (with maximum confidence ratings) relied solely on the minimal learning rate.

Predictive learning followed the standard Rescorla-Wagner learning model (56):

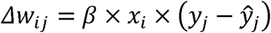

Here, the model updated the weight w_ij_ based on the error between the actual feedback y_j_ and the model’s prediction 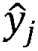.

After the learning phase, the model completed a final assessment, replicating the procedure used in Phase 4 of the human task. The activations in the output layer were normalized as follows:

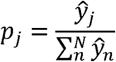

with the activation level (p_j_) of the correct Swahili unit serving as an indicator of the model’s accuracy for each trial.

### Model fitting

Since the main goal was to replicate human responses in Phase 4, the model’s final assessment results were used to fit the human final assessment data, which determined the optimal model parameters. The optimization was conducted by maximizing the log-likelihood function for each participant’s data. Parameters α and β were allowed to range from 0 to 1, while *s* and *b* were allowed to range from 0 to 10. Additionally, the sum of α and β was constrained to 1, ensuring the relative contribution of initial learning versus Hebbian learning (and/or predictive learning) was accounted for.

### Model comparison

Importantly, we developed seven models based on all possible combinations of the three learning components mentioned earlier (2³ – 1 = 7): initial, Hebbian, and predictive learning. An additional model (Model 8) incorporated initial learning and retrieval effort learning. The weighted Akaike information criterion (wAIC; higher means better) was employed to compare the model fits (57).

### fMRI data acquisition

fMRI data were collected with a 3T Prisma scanner system (Siemens), with a 64-channel head coil. A 3D high-resolution anatomic image was obtained using a T1-weighted MPRAGE sequence [flip angle (FA) = 9°; matrix size = 176 × 256 × 256; interpolated voxel size = 1.0 × 1.0 × 1.0mm]. In addition, we acquired a field map per participant, to correct for magnetic field inhomogeneities (TR= 528ms; TE = 7.38ms; FA= 60°). Whole-brain functional images were acquired using a T2p-weighted EPI sequence (TR = 1780ms; TE = 27ms; FA= 66°; matrix size = 54 × 64 × 64; voxel size = 2.5 × 2.5 × 2.5 mm).

### fMRI data analysis

#### Preprocessing

The fMRI data analysis was conducted using SPM 12 and MATLAB. After converting the raw DICOM data to BIDS format, the first five functional images for each participant were discarded to avoid transient spin saturation effects. The remaining functional images underwent slice timing correction, realignment, normalization, and smoothing. The structural image was co-registered with the mean realigned functional image. Event-related regressors (delta functions) were convolved with the canonical Hemodynamic Response Function (HRF) in the General Linear Model (GLM) to model the BOLD signal.

#### GLM for Test vs Study

In Test-GLM1, we modeled the choice onsets (correct test, incorrect test, and study) and feedback onsets (correct test, incorrect test, and study) as six main regressors. Here and in all other GLMs, fixation onsets and confidence onsets were always modeled as additional regressors. Beta maps from these regressors generate individual contrasts for group-level analysis (voxel level; False Discovery Rate, FDR corrected *p* < 0.05). First, we contrasted the beta maps of Test vs. Study (0.5 × correct test + 0.5 × incorrect test − 1 × study), Correct test vs. Study (1 × correct test − 1 × study), and Incorrect test vs. Study (1 × incorrect test − 1 × study) during feedback onsets to identify significantly stronger brain activation in test, correct test, and incorrect test conditions (compared to study). Second, the same neural contrasts were applied for choice onsets. Test-GLM2 modeled same regressors as Test-GLM1 but added trial-level confidence ratings as the parametric modulator of (testing) choice onset regressors.

#### GLM for prediction error

We constructed two general linear models to examine the neural correlates of prediction errors. In the PE-GLM1, feedback onsets were modeled as a single main regressor, with trial-level (mean-centered) prediction errors and feedback valence jointly included as parametric modulators of feedback onsets. Individual beta maps corresponding to the prediction error modulator were submitted to group-level analysis (voxel level; FDR corrected *p* < 0.05). Brain regions showing positive activation in the group-level analysis were interpreted as being positively modulated by prediction error. Neural responses associated with the feedback valence modulator were examined using the same procedure.

The overlapping active brain regions between the testing and prediction error were defined as ROIs for PE-GLM2. In this model, feedback onsets associated with different prediction error levels in the testing condition and in the studying condition (only zero prediction error in this case) were modeled as separate regressors. To avoid creating regressors for nearly every trial, prediction errors were rounded to one decimal place. For each participant, we extracted the mean beta values within each ROI for each prediction error regressor. The relationship between prediction errors and mean beta values (of each ROI) across all participants was then visualized in Figure 4.

#### Back sorting analysis

In the first and second back sorting GLMs (Bsort-GLM1 and Bsort-GLM2), feedback onsets followed by correct Phase 4 assessment and those followed by incorrect Phase 4 assessment were modeled as two main regressors. Individual contrast maps (Correct - Incorrect) were submitted to group-level analysis (cluster level; cluster-forming threshold at *p* < 0.005; cluster-level significance at FDR-corrected *p* < 0.05). Bsort-GLM3 was constructed in the same manner for choice onsets.

## Acknowledgments

We thank Jonas Simoens, Ruth Krebs, and Filip Van Opstal for useful discussions about the topic of this study. H.C. received support from China Scholarship Council [CSC202206990008]. C.B.C. discloses support for the publication of this work from Centro Nacional de Inteligencia Artificial [FB210017]. The funders had no role in study design, data collection and analysis, decision to publish or preparation of the manuscript.

## Footnotes

* Final assessment accuracy was coded as 1 for correct responses in both final assessments, 0.5 for one correct response, and 0 for no correct response.

† Statistical results remained unchanged when only a random intercept was added.

‡ A score of 0.5 means a lucky guess rather than intermediate knowledge between know and not know.

§ Another binary strategy encoded final assessment accuracy as 1 for correct responses in both final assessments, and 0 otherwise, which led to the same statistical results.

## References

1. K. B. McDermott, Practicing Retrieval Facilitates Learning. Annual Review of Psychology 72, 609–633 (2021).

2. H. L. Roediger, A. C. Butler, The critical role of retrieval practice in long-term retention. Trends in Cognitive Sciences 15, 20–27 (2011).

3. H. L. Roediger, J. D. Karpicke, Test-enhanced learning: Taking memory tests improves long-term retention. Psychological science 17, 249–255 (2006).

4. B. Jonsson, C. Wiklund-Hörnqvist, T. Stenlund, M. Andersson, L. Nyberg, A learning method for all: The testing effect is independent of cognitive ability. Journal of Educational Psychology 113, 972–985 (2021).

5. T. R. Jonker, H. Dimsdale-Zucker, M. Ritchey, A. Clarke, C. Ranganath, Neural reactivation in parietal cortex enhances memory for episodically linked information. Proceedings of the National Academy of Sciences 115, 11084–11089 (2018).

6. J. D. Karpicke, H. L. Roediger, Expanding retrieval practice promotes short-term retention, but equally spaced retrieval enhances long-term retention. Journal of Experimental Psychology: Learning, Memory, and Cognition 33, 704–719 (2007).

7. M. A. Pyc, K. A. Rawson, Testing the retrieval effort hypothesis: Does greater difficulty correctly recalling information lead to higher levels of memory? Journal of Memory and Language 60, 437–447 (2009).

8. S. K. Carpenter, E. L. DeLosh, Impoverished cue support enhances subsequent retention: Support for the elaborative retrieval explanation of the testing effect. Memory & cognition 34, 268–276 (2006).

9. S. K. Carpenter, Cue strength as a moderator of the testing effect: The benefits of elaborative retrieval. Journal of Experimental Psychology: Learning, Memory, and Cognition 35, 1563–1569 (2009).

10. H. Chen, C. Hauspie, K. Ergo, C. Buc Calderon, T. Verguts, Predictive learning as the basis of the testing effect. Communications Psychology 3, 18 (2025).

11. Y. Zheng, X. L. Liu, S. Nishiyama, C. Ranganath, R. C. O’Reilly, Correcting the hebbian mistake: Toward a fully error-driven hippocampus. PLOS Computational Biology 18, e1010589 (2022).

12. X. L. Liu, R. C. O’Reilly, C. Ranganath, “Chapter Four - Effects of retrieval practice on tested and untested information: Cortico-hippocampal interactions and error-driven learning” in Psychology of Learning and Motivation, K. D. Federmeier, L. Sahakyan, Eds. (Academic Press, 2021), vol. 75, pp. 125–155.

13. R. Sutton, A. Barto, Reinforcement learning: an introduction (MIT, Cambridge, MA, 2018).

14. E. De Loof et al., Signed reward prediction errors drive declarative learning. PLoS One 13, e0189212 (2018).

15. N. Rouhani, K. A. Norman, Y. Niv, Dissociable effects of surprising rewards on learning and memory. Journal of Experimental Psychology: Learning, Memory, and Cognition 44, 1430 (2018).

16. A. Greve, E. Cooper, A. Kaula, M. C. Anderson, R. Henson, Does prediction error drive one-shot declarative learning? Journal of Memory and Language 94, 149–165 (2017).

17. R. Daniel, S. Pollmann, Striatal activations signal prediction errors on confidence in the absence of external feedback. NeuroImage 59, 3457–3467 (2012).

18. T. Balsdon, M. A. Pisauro, M. G. Philiastides, Distinct basal ganglia contributions to learning from implicit and explicit value signals in perceptual decision-making. Nature Communications 15, 5317 (2024).

19. T. D. Satterthwaite et al., Being right is its own reward: Load and performance related ventral striatum activation to correct responses during a working memory task in youth. NeuroImage 61, 723–729 (2012).

20. C. B. Holroyd et al., Dorsal anterior cingulate cortex shows fMRI response to internal and external error signals. Nature neuroscience 7, 497–498 (2004).

21. N. Eshel, J. Tian, M. Bukwich, N. Uchida, Dopamine neurons share common response function for reward prediction error. Nature Neuroscience 19, 479–486 (2016).

22. P. Montague, P. Dayan, T. Sejnowski, A framework for mesencephalic dopamine systems based on predictive Hebbian learning. The Journal of Neuroscience 16, 1936–1947 (1996).

23. W. Schultz, P. Dayan, P. R. Montague, A Neural Substrate of Prediction and Reward. Science 275, 1593–1599 (1997).

24. K. D’Ardenne, S. M. McClure, L. E. Nystrom, J. D. Cohen, BOLD Responses Reflecting Dopaminergic Signals in the Human Ventral Tegmental Area. Science 319, 1264–1267 (2008).

25. C. B. Calderon et al., Signed Reward Prediction Errors in the Ventral Striatum Drive Episodic Memory. J Neurosci 41, 1716–1726 (2021).

26. L. K. Fazio, E. J. Marsh, Correcting false memories. Psychological science 21, 801–803 (2010).

27. J. Metcalfe, B. Finn, People’s hypercorrection of high-confidence errors: Did they know it all along? Journal of Experimental Psychology: Learning, Memory, and Cognition 37, 437 (2011).

28. P. Ripollés et al., The Role of Reward in Word Learning and Its Implications for Language Acquisition. Current Biology 24, 2606–2611 (2014).

29. T. Verguts, Introduction to Modeling Cognitive Processes (MIT Press, 2022).

30. D. F. Wood, Problem based learning. BMJ 326, 328–330 (2003).

31. M. Prince, Does Active Learning Work? A Review of the Research. Journal of Engineering Education 93, 223–231 (2004).

32. C. Yang, L. Luo, M. A. Vadillo, R. Yu, D. R. Shanks, Testing (quizzing) boosts classroom learning: A systematic and meta-analytic review. Psychological Bulletin 147, 399–435 (2021).

33. J. L. McClelland, B. L. McNaughton, R. C. O’Reilly, Why there are complementary learning systems in the hippocampus and neocortex: insights from the successes and failures of connectionist models of learning and memory. Psychol Rev 102, 419–457 (1995).

34. A. Pine, N. Sadeh, A. Ben-Yakov, Y. Dudai, A. Mendelsohn, Knowledge acquisition is governed by striatal prediction errors. Nature communications 9, 1–14 (2018).

35. B. Butterfield, J. A. Mangels, Neural correlates of error detection and correction in a semantic retrieval task. Cognitive Brain Research 17, 793–817 (2003).

36. J. Metcalfe, B. Butterfield, C. Habeck, Y. Stern, Neural correlates of people’s hypercorrection of their false beliefs. Journal of Cognitive Neuroscience 24, 1571–1583 (2012).

37. B. Butterfield, J. Metcalfe, Errors committed with high confidence are hypercorrected. Journal of Experimental Psychology: Learning, Memory, and Cognition 27, 1491 (2001).

38. T. Seabrooke, C. J. Mitchell, A. J. Wills, A. B. Inkster, T. J. Hollins, The benefits of impossible tests: Assessing the role of error-correction in the pretesting effect. Memory & Cognition 50, 296–311 (2022).

39. M. Watabe-Uchida, N. Eshel, N. Uchida, Neural Circuitry of Reward Prediction Error. Annu Rev Neurosci 40, 373–394 (2017).

40. W. Schultz, Reward prediction error. Current Biology 27, R369–R371 (2017).

41. S. J. Sara, The locus coeruleus and noradrenergic modulation of cognition. Nature Reviews Neuroscience 10, 211–223 (2009).

42. P. T. Ohara et al., Dopaminergic input to GABAergic neurons in the rostral agranular insular cortex of the rat. Journal of Neurocytology 32, 131–141 (2003).

43. M. G. Kutlu et al., Role of insular cortex D₁ and D₂ dopamine receptors in nicotine self-administration in rats. Behav Brain Res 256, 273–278 (2013).

44. N. Gogolla, The insular cortex. Current Biology 27, R580–R586 (2017).

45. E. Hernández-Ortiz, J. Luis-Islas, F. Tecuapetla, R. Gutierrez, F. Bermúdez-Rattoni, Top-down circuitry from the anterior insular cortex to VTA dopamine neurons modulates reward-related memory. Cell Reports 42, 113365 (2023).

46. C. W. Hoy et al., Asymmetric coding of reward prediction errors in human insula and dorsomedial prefrontal cortex. Nature Communications 14, 8520 (2023).

47. J. Metcalfe, M. Vuorre, E. Towner, T. S. Eich, Curiosity: The effects of feedback and confidence on the desire to know. J Exp Psychol Gen 152, 464–482 (2022).

48. K. Ergo, E. De Loof, T. Verguts, Reward Prediction Error and Declarative Memory. Trends in Cognitive Sciences 24, 388–397 (2020).

49. Juliet Y. Davidow, K. Foerde, A. Galván, D. Shohamy, An Upside to Reward Sensitivity: The Hippocampus Supports Enhanced Reinforcement Learning in Adolescence. Neuron 92, 93–99 (2016).

50. A. I. Jang, M. R. Nassar, D. G. Dillon, M. J. Frank, Positive reward prediction errors during decision-making strengthen memory encoding. Nature human behaviour 3, 719–732 (2019).

51. K. Ergo, E. De Loof, C. Janssens, T. Verguts, Oscillatory signatures of reward prediction errors in declarative learning. Neuroimage 186, 137–145 (2019).

52. N. Kornell, M. J. Hays, R. A. Bjork, Unsuccessful retrieval attempts enhance subsequent learning. Journal of Experimental Psychology: Learning, Memory, and Cognition 35, 989–998 (2009).

53. N. Kornell, P. J. Klein, K. A. Rawson, Retrieval attempts enhance learning, but retrieval success (versus failure) does not matter. Journal of Experimental Psychology: Learning, Memory, and Cognition 41, 283 (2015).

54. S. Bertsch, B. J. Pesta, R. Wiscott, M. A. McDaniel, The generation effect: A meta-analytic review. Memory & Cognition 35, 201–210 (2007).

55. S. S. H. Wong, S. W. H. Lim, The derring effect: Deliberate errors enhance learning. Journal of Experimental Psychology: General 151, 25–40 (2022).

56. R. A. Rescorla, A. R. Wagner, “A theory of Pavlovian conditioning: variations in the effectiveness of reinforcement and nonreinforcement” in Classical Conditioning II: Current Research and Theory, A. H. Blake, W. F. Prokasy, Eds. (Appleton-Century-Croft, 1972), pp. 64–99.

57. E.-J. Wagenmakers, S. Farrell, AIC model selection using Akaike weights. Psychonomic bulletin & review 11, 192–196 (2004).

