## Supplementary Information for "Neural and computational evidence for a predictive learning account of the testing effect"

*Haopeng Chen

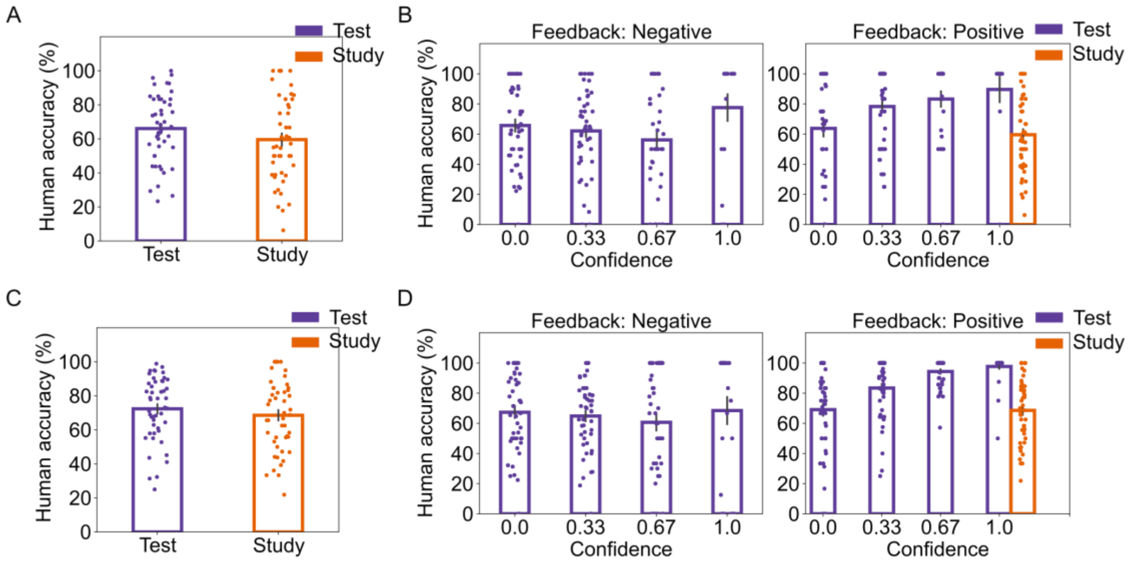


**Figure S1. The behavioural patterns after removing correct or high-confidence word pairs in Phase 2. A-B.** The testing effect is still significant after removing correct word pairs in Phase 2 ($\chi^2$(1, N = 48) = 6.563, *p* = .010, *β* ± *95% CI* = .063 ± .049, $R^2$ = .004). **C-D.** The testing effect is still significant after removing high confidence (confidence = 1) word pairs in Phase 2 ($\chi^2$(1, N = 48) = 6.149, *p* = .013, *β* ± *95% CI* = .039 ± .032, $R^2$ = .002). Individual data points are represented by the dots; The error bars reflect the standard errors; Confidence ratings were from Phase 3; $p < 0.05$*. *N* (subjects) = 48.

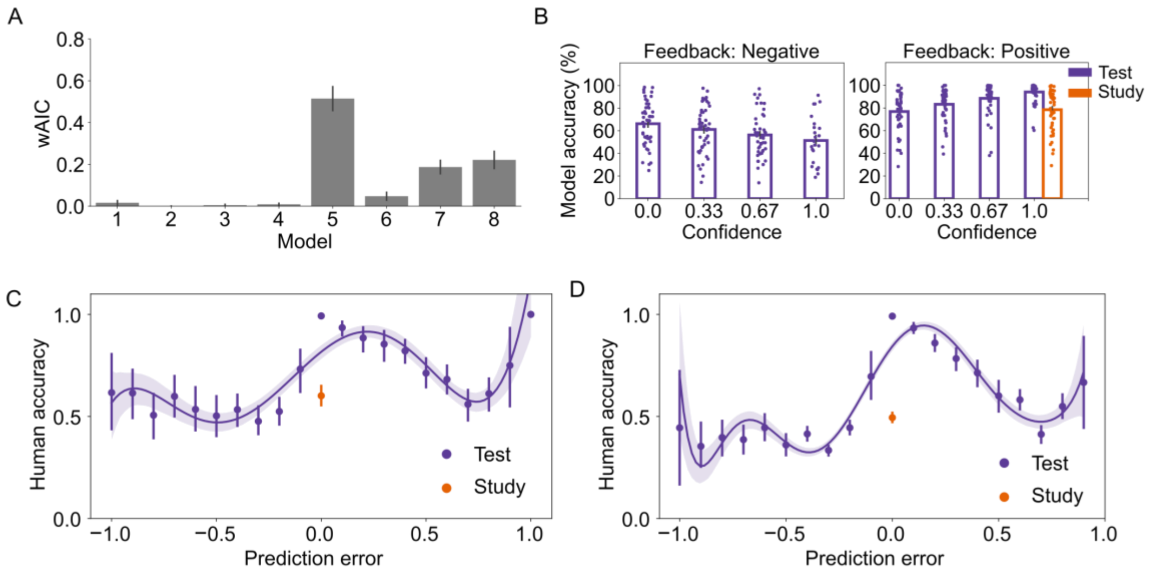


**Figure S2. Additional modeling results**. **A.** Model comparison with wAIC (higher means better) suggests that the model with both initial learning and predictive learning (Model 5) is still the best model. **B.** The model with both initial learning and Retrieval effort (Hebbian) learning could not replicate the testing effect in human data. **C-D.** In a previous study using the same experimental design and computational model (1)**,** prediction errors also influence immediate (10 minutes after Phase 3) and delayed (24 hours after Phase 3) final assessment accuracy in a W-shaped pattern. Individual data points are represented by the dots; The error bars reflect the standard errors; Confidence ratings were from Phase 3; $p$ < 0.05*. $N$ (subjects) = 48.

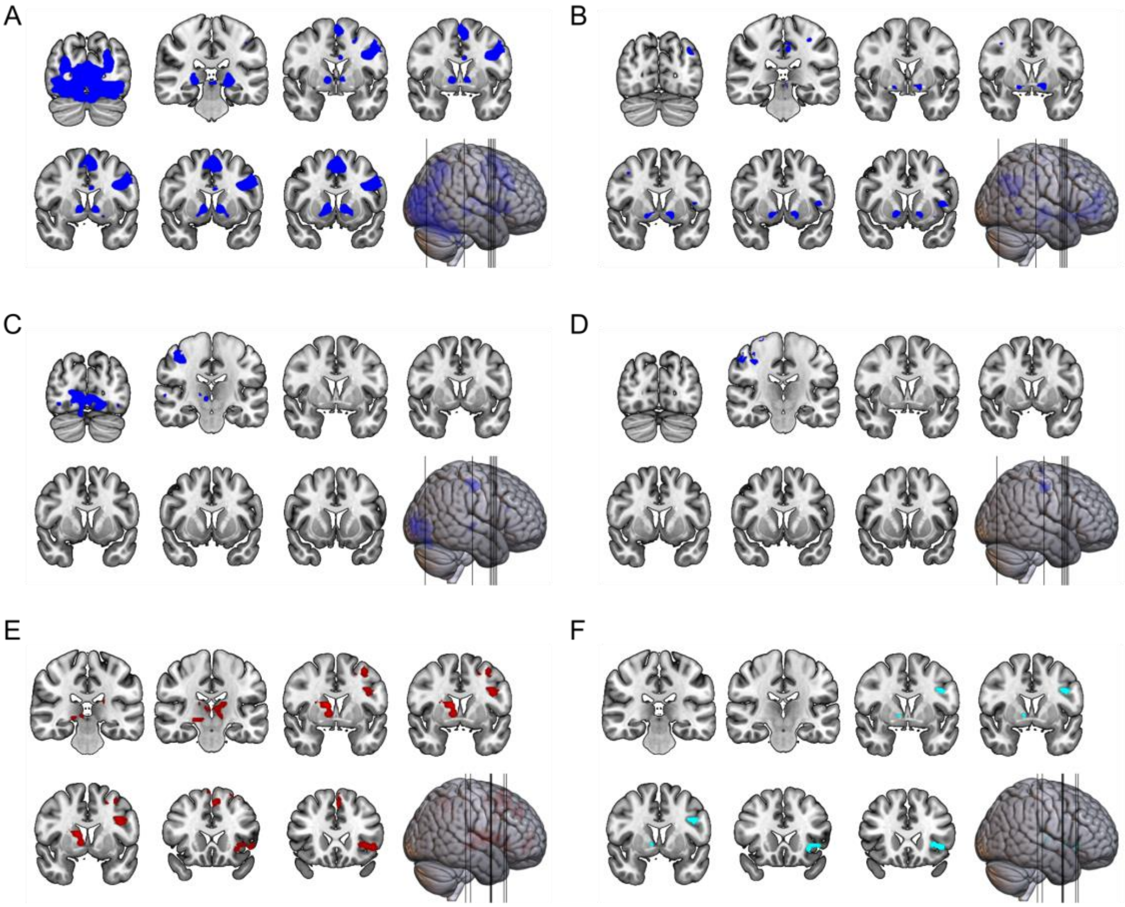


**Figure S3. Additional fMRI results**. **A.** Test (vs Study) elicited stronger neural activations in the VS and visual cortex during choice onsets. **B.** The VS activation during choice onsets was positively modulated by confidence ratings. **C-D.** Test (vs Study) did not elicit any significant VS activation during two fixation periods (C: earlier, D: later) between choice onsets and feedback onsets. **E-F.** In an additional back-sorting analysis that included all incorrect trials and a broader set of correct trials (prediction error ≥ 0.5) in Phase 3, we still found VS and insula activations (E), which overlapped with those elicited by testing and prediction errors (F). *N* (subjects) = 48.

### SI References


1. H. Chen, C. Hauspie, K. Ergo, C. Buc Calderon, T. Verguts, Predictive learning as the basis of the testing effect. *Communications Psychology* **3**, 18 (2025).
